# Narcolepsy type 1 patients have abnormal brain activation to neutral-rated movies in humor-paradigm

**DOI:** 10.1101/377499

**Authors:** Hilde T. Juvodden, Dag Alnæs, Martina J. Lund, Espen Dietrichs, Per M. Thorsby, Lars T. Westlye, Stine Knudsen

## Abstract

Narcolepsy type 1 is a neurological sleep disorder mainly characterized by excessive daytime sleepiness, fragmented night sleep, and cataplexy (muscle atonia triggered by emotions). To characterize brain activation patterns in response to neutral-rated and fun-rated movies in narcolepsy type 1 we performed functional magnetic resonance imaging during a paradigm consisting of 30 short movies (25/30 with a humorous punchline; 5/30 without a humorous punchline (but with similar build-up/anticipation)) that the participants rated based on their humor experience. We included 41 narcolepsy type 1 patients (31 females, mean age 23.6 years, 38/41 H1N1-vaccinated, 41/41 HLA-DQB1*06:02-positive, 40/40 hypocretin-deficient) and 44 first-degree relatives (24 females, mean age 19.6 years, 30/44 H1N1-vaccinated, 27/44 *HLA-DQB1*06:02-positive*) as controls. Group-level inferences were made using permutation testing.

Permutation testing revealed no significant differences in the average ratings of patients and controls. Functional magnetic resonance imaging analysis revealed that both groups showed higher activations in response to fun-rated movies in several brain regions associated with humor processing, with no significant group differences. In contrast, patients showed significantly higher activation compared to controls during neutral-rated movies; including bilaterally in the thalamus, pallidum, putamen, amygdala, hippocampus, middle temporal gyrus, cerebellum, brainstem and in the left precuneus, supramarginal gyrus and caudate.

The presence of a humorous punchline in a neutral-rated movie is important since we found no brain overactivation for narcolepsy type 1 patients for movies without a humorous punchline (89.0% neutral-rated) compared with controls.

Further, a comparison between fun-rated and neutral-rated movies revealed a pattern of higher activation during fun-rated movies in controls, patients showed no significant differentiation between these states. Group analyses revealed significantly stronger differentiation between fun-rated and neutral-rated movies in controls compared with patients, including bilaterally in the inferior frontal gyrus, thalamus, putamen, precentral gyrus, lingual gyrus, supramarginal gyrus, occipital areas, temporal areas, cerebellum and in the right hippocampus, postcentral gyrus, pallidum and insula.

In conclusion, during neutral-rated movies, narcolepsy type 1 patients showed significantly higher activation in several cortical and subcortical regions previously implicated in humor and REM sleep, including the thalamus and basal ganglia. The relative lack of differentiation between neutral-rated and fun-rated movies in narcolepsy type 1 patients might represent insight into the mechanisms associated with cataplexy, in which a long-lasting hypervigilant state could represent risk (hypersensitivity to potential humorous stimuli) for the narcolepsy type 1 patients, which seem to have a lower threshold for activating the humor response, even during neutral-rated movies.

## Introduction

Narcolepsy type 1 is a disabling, chronic neurological sleep disorder primarily characterized by excessive daytime sleepiness, cataplexy and sleep-onset REM-periods. Patients also experience fragmented night sleep, hypnagogic/hypnopompic hallucinations and sleep paralysis (American Academy of Sleep Medicine(AASM), 2014; Dauvilliers *et al*., 2014; Kornum *et al*., 2017). Narcolepsy type 1 patients are hypocretin (also called orexin)-deficient due to loss of hypocretin producing hypothalamic neurons (Peyron *et al*., 2000; Thannickal *et al*., 2000), with projections widely distributed in the brain (Peyron *et al*., 1998; Li *et al*., 2014; Mahler *et al*., 2014). Hypocretin 1 and 2 (Orexin-A and B) are central regulators of sleep-wake and muscle tonus (Saper *et al*., 2010; Saper, 2013). Autoimmune destruction of hypocretin-producing neurons is hypothesized to be the pathogenesis of narcolepsy type 1 (Kornum *et al*., 2017), which is further supported by the >10-fold increase of H1N1-vaccine related narcolepsy type 1 cases occurring after the H1N1 flu vaccination campaigns with Pandemrix^®^ in 2009/2010 in several European countries, including Norway (Heier *et al*., 2013).

Cataplexy, emotionally triggered involuntary muscle weakness or paralysis during wakefulness, is a defining clinical feature of narcolepsy type 1 (Dauvilliers *et al*., 2014). Episodes can typically last from several seconds to several minutes with retained consciousness, usually triggered by strong, positive emotions (for example thinking of, hearing or telling a joke), although various emotions can be triggers. Cataplectic episodes can also be reported by patients to be unrelated to emotions and have no identifiable trigger (Dauvilliers *et al*., 2014; Kornum *et al*., 2017). The frequency of cataplexy episodes varies greatly; while some patients seldom have episodes, others can have more than 20 per day. There is also variation in the degree of muscle atonia involvement in cataplexy, from partial slight hypotonia to a complete inability to move (Kornum *et al*., 2017). It has also been reported (Overeem *et al*., 2011) that some patients can feel cataplexy attacks coming on with warning signs and learn to avoid cataplexy by avoiding triggering situations (Dauvilliers *et al*., 2007; Overeem *et al*., 2011).

Three previous functional MRI studies used humorous pictures (Reiss *et al*., 2008; Schwartz *et al*., 2008) or movies (Meletti *et al*., 2015) to study humor-processing and cataplexy in sporadic (non-vaccinated) narcolepsy type 1 patients. These studies involved a relatively small number of patients (n = 10-21). The largest study (Meletti *et al*., 2015) acquired with simultaneous EEG, attempted to elicit cataplexy in patients with a naturalistic paradigm in which the humorous movies were selected in accordance with each patient’s humor preferences, while the other two studies (Reiss *et al*., 2008; Schwartz *et al*., 2008) primarily assessed humor processing in narcolepsy type 1 patients.

Schwarz et al. found lower activation in several areas including the hypothalamus and higher activation in several areas including the amygdala for humorous pictures for narcolepsy type 1 patients compared with controls. Reiss et al. found higher activations for narcolepsy type 1 patients compared with controls when looking at funny cartoons compared to non-funny cartoons in several regions, including; hypothalamus, ventral striatum and right inferior frontal gyrus. Meletti et al. found that laughter was associated with higher activation bilaterally in anterior cingulate gyrus and the motor/premotor cortex, and that cataplexy was associated with higher activation in several areas, including; the amygdala, anterior insula, ventromedial prefrontal cortex, nucleus accumbens, locus coeruleus and the anteromedial pons.

The mechanism underlying cataplexy has not been established (Kornum *et al*., 2017). One theory suggests that cataplexy is a form of tonic immobility that can be seen in some animals (Overeem *et al*., 2002), while another states that cataplexy represents dissociated REM sleep that appears while the person is awake (Dauvilliers *et al*., 2014; Kornum *et al*., 2017). Due to cataplexy possibly representing dissociated REM sleep we were particularly interested in regions that have been implicated in REM sleep and cataplexy/humor (thalamus, amygdala and basal ganglia). Previous studies using functional MRI (Wehrle *et al*., 2005; Wehrle *et al*., 2007; Miyauchi *et al*., 2009) and PET (Maquet *et al*., 1996; Braun *et al*., 1997; Maquet and Franck, 1997; Buchsbaum *et al*., 2001) in healthy individuals with polysomnographic monitoring have revealed higher thalamus activation during REM sleep. Further, the thalamus has been shown to have higher activity in response to humor processing (Vrticka *et al*., 2013), with higher activation in response to cartoons compared with neutral pictures (Mobbs *et al*., 2003; Kohn *et al*., 2011) and during humor-induced smiling (Wild *et al*., 2006).

The amygdala has been reliably associated with humor appreciation in humans (Vrticka *et al*., 2013). PET (Maquet *et al*., 1996; Maquet and Franck, 1997; Nofzinger *et al*., 1997) and functional MRI (Miyauchi *et al*., 2009) studies have also shown amygdala activation during REM sleep in healthy individuals. Increased basal ganglion activation has also been linked to REM sleep (Braun *et al*., 1997; Wehrle *et al*., 2005; Miyauchi *et al*., 2009) and humor (Phan *et al*., 2002; Goldin *et al*., 2005; Wild *et al*., 2006).

The conflicting results from the previous functional MRI studies of sporadic narcolepsy (Reiss *et al*., 2008; Schwartz *et al*., 2008; Meletti *et al*., 2015) are probably due primarily to a combination of low power, which inevitably challenges the reliability, and different behavioral paradigms. Our study focuses on humor processing, as partially explored by (Reiss *et al*., 2008; Schwartz *et al*., 2008), but using a larger sample size. The previous studies included ratings of pictures, but only reported results of neutral-rated pictures in relation to fun-rated pictures. Here, we assess brain activation patterns during fun-rated and neutral-rated short movies, including movies with and without a humorous punchline.

To test the hypothesis of abnormal humor processing in narcolepsy type 1, we compared functional MRI-based brain activation patterns during presentations of short movies in 41 narcolepsy type 1 patients (31 females, mean age 23.6 years, 41/41 *HLA-DQB1*06:02-* postive, 38/41 H1N1-vaccinated and 40/40 hypocretin deficient (1 patient had not yet performed this measure) and a control group of 44 first-degree relatives of narcolepsy type 1 patients (24 females, 19.6 years, 27/44 *HLA-DQB1*0602*-positive, 30/44 H1N1-vaccinated). We tested for group differences across the whole brain and corrected for non-independence due to familiarity and for multiple comparisons by permutation testing.

## Materials and methods

### Participants

Table 1 summarizes the demographic and clinical information about the two groups. We have previously reported on 40 patients and 44 controls considered in the current study (Juvodden *et al*., 2018). Participants were recruited from those who were referred for narcolepsy family disease education and counseling courses at the Norwegian Centre of Expertise for Neurodevelopmental Disorders and Hypersomnias (NevSom) during the inclusion period from June 2015 to April 2017. 41 patients with narcolepsy type 1, with disease onset after the H1N1 vaccination in 2009/2010, and 44 first-degree relatives of narcolepsy type 1 patients were included consecutively. Not all first-degree relatives of narcolepsy type 1 patients were 1^st^ degree relatives of patients included in this study, as sometimes patients were excluded, but all these excluded patients had a verified narcolepsy type 1 diagnosis. Subsequently, the disease onset was changed to being before H1N1 vaccination for three narcolepsy type 1 patients after a thorough evaluation of their medical history and records (3/3 with typical narcolepsy type 1 phenotypes, being *HLA-DQB1*06:02-positive*, hypocretin-deficient with cataplexy, and were therefore kept in the study). Written informed consent was provided by all participants before inclusion, and the Norwegian regional committees for medical and health research ethics (REK) approved our study. The official Norwegian Immunization Registry (SYSVAK) was used to obtain H1N1-vaccination status of patients and first-degree relatives. Pandemrix^®^ was the only vaccine used for H1N1-vaccination in Norway. Two patients, not registered in SYSVAK, who reported having been H1N1-vaccinated in their workplace were also included in the H1N1-vaccinated group.

**Table 1.** Demographic and clinical data.

|  | Narcolepsy type 1<br>(n=41) | First-degree relatives<br>(n=44) |
| --- | --- | --- |
| Gender (female), n (%) | 31 (75.6) | 24 (54.5) |
| Age (years), Mean ± SD | 23.6 ± 11.4 | 19.6 ± 8.4 |
| Age at disease onset (years), mean ± SD | 17.7 ± 10.9 | N/A |
| Disease duration (years) mean ± SD | 5.9 ± 1.5 | N/A |
| H1N1-vaccinated, n (%) | 38 (92.7) | 30 (68.2) |
| Cataplexy, n (%) | 39 (95.1) | 6 (13.6)* |
| *HLA-DQB1* *06:02-positivity, n (%) | 41 (100) | 27 (61.4) |
| CSF hypocretin-1 ≤ 1/3 of level in<br>normal population | 40/40<br>(1-N/A) | N/A |
| Hypnagogic hallucinations, n (%) | 35 (85.4) | 9 (20.5) |
| Sleep paralysis, n (%) | 29 (70.7) | 8 (18.2) |

N/A: not available, SD: standard deviation. Disease onset was reclassified to being before the
H1N1-vaccinations for three narcolepsy type 1 patients after a thorough evaluation of their medical history and records (3/3 were typical narcolepsy type 1 phenotypes, hypocretin-deficient, *HLA-DQB1*\*06:02-positive with cataplexy, and so were retained in the study).
\*First-degree relatives with signs of cataplexy all experienced it rarely, but with triggers known to elicit cataplexy, like laughter, fun, excitement and surprise

Fourteen days before their inclusion, all patients had ceased all narcolepsy medication, except for one patient who, due to severe cataplexy, was without narcolepsy medication for only 7 days. Exclusion criteria for patients and first-degree relatives were severe neurological, psychiatric or somatic disorders, previous head injury with loss of consciousness for 10 minutes or 30 minutes amnesia, metallic implants, excessive movement during the MRI-scanning, and neuroradiological findings requiring clinical follow-up. Since narcolepsy type 1 is associated with an increased number of comorbidities (Kornum *et al*., 2017) we performed the analysis in the full sample including the comorbidities listed below (for patients and first-degree relatives) and in a reduced sample, from which we excluded patients and first-degree relatives with the comorbidities. In the reduced sample we also excluded participants who had to re-watch the movies (mainly due to drowsiness/falling asleep during the first viewing; *n* = 8) or watch the movies in black and white (due to a technical problem; *n* = 1), or without sound (due to a human error; *n* = 2), and first-degree relatives who had experienced cataplexy-like episodes (*n* = 6).

In the full sample analysis, the following comorbidities were present in the narcolepsy type 1 patients: Asperger syndrome (*n* = 2), attention deficit hyperactivity disorder (ADHD, *n* = 1), migraine (*n* = 4), Tourette syndrome (*n* = 1), anxiety (*n* = 1), depression (*n* = 1), prematurity without severe long-term complications (*n* = 1), kidney disease (*n* = 1), type 2 diabetes (*n* = 1) and hypothyroidism (*n* = 2). We also accepted the following morbidities in the first-degree relatives group for the full sample analysis: attention deficit disorder (ADD, *n* = 2), migraine (*n* = 6), dyslexia (*n* = 4), anxiety (*n* = 1), prematurity without severe long-term complications (*n* = 2) and bipolar type 2 disorder (*n* = 1). Some patients and first-degree relatives had more than one comorbidity.

All patients (41/41) and 61.4% (27/44) of the first-degree relatives were *HLA-DQB1*06:02-* positive. All patients whose hypocretin level was measured (*n* = 40) were hypocretin-deficient (CSF hypocretin-1 level < 110 pg/ml or < 1/3 of the normal mean, as previously reported (Heier *et al*., 2007; Knudsen *et al*., 2010)), while one HLA-DQB1*0602-positive patient with typical cataplexy had not yet had this measured. 92.7% (38/41) of all patients and 68.2% (30/44) of first-degree relatives were H1N1-vaccinated. 95.1% (39/41) of all patients reported having had cataplexy episodes. 13.6% (6/44) of the first-degree relatives reported signs of muscle weakness, although they all experienced this rarely, triggered by emotions known to elicit cataplexy: laughter, fun/excitement and surprise. 85.4% (35/41) of patients reported hypnagogic hallucinations and 70.7% (29/41) experienced sleep paralysis. 20.5% (9/44) of first-degree relatives also experienced hypnagogic hallucinations and 18.2% (8/44) had experienced sleep paralysis.

### Narcolepsy diagnosis

International Classification of Sleep Disorders (ICSD)-3 criteria were used to establish the narcolepsy type 1 diagnoses (American Academy of Sleep Medicine(AASM), 2014) given by the experienced neurologist and sleep medicine expert Stine Knudsen.

Patients and first-degree relatives completed clinical consultations, a neurological examination, routine blood samples, actigraphy, polysomnography, the multiple sleep latency test (MSLT) and HLA typing. Participants also took part in semi-structured interviews about narcolepsy and sleep disorders, including a Norwegian translation of the Stanford Sleep Questionnaire (Anic-Labat *et al*., 1999). Measurements of CSF hypocretin-1 levels in patients were also obtained (Phoenix Pharmaceutical St. Joseph, MO, USA), slightly modified, and analyzed at the Hormone Laboratory, Oslo University Hospital) (Heier *et al*., 2007; Knudsen *et al*., 2010).

After taking into consideration clinical evaluation, polysomnography, MSLT and hypocretin measures, all patients fulfilled the ICSD-3 criteria for narcolepsy. No first-degree relatives met the ICSD-3 criteria for narcolepsy.

### Polysomnography recordings

10-14 days of actigraphy (Philips Actiwatch, Respironics Inc., Murrysville, PA, USA) preceded all polysomnography recordings. International Classification of Sleep Disorders (ICSD)-3 criteria (American Academy of Sleep Medicine(AASM), 2014) were used to evaluate all participants with polysomnography and MSLT. The SOMNOmedics system (SOMNOmedics GmbH, Randersacker, Germany) was used to obtain polysomnography recordings with the F3-A2, C3-A2, O1-A2, F4-A1, C4-A1 and O2-A1 electrodes, in addition to vertical and horizontal electro-oculography, surface EMG of the submentalis and tibialis anterior muscles, electrocardiography, nasal air flow, thoracic respiratory effort and oxygen saturation. EMG impedance was kept below 10kΩ (preferably 5Ω). AASM criteria (American Academy of Sleep Medicine(AASM), 2014) were applied to the sleep scoring.

### Functional MRI paradigm

Participants watched 30 movies by looking through a mirror attached to a 32-channel headcoil to view an MR-compatible LCD screen (NNL LCD Monitor^®^, NordicNeuroLab, Bergen, Norway) behind the scanner. Stimuli were presented using E-Prime 2.0 software (Psychology Software Tools, Pittsburgh, PA). 25/30 movies included a humorous punchline, while the humorous punchline had been edited out of the remaining five. Movies with and without a humorous punchline varied in duration from 10-20 s (mean 14.4 ± 3.5 s) and from 10-15 s (mean 12.0 ± 2.3 s), respectively. Both types of movie had similar build-up/anticipation that something funny might happen.

Participants rated all movies (with or without a humorous punchline) using an MR-compatible subject response collection system (ResponseGrip^®^, NordicNeuroLab, Bergen, Norway) to choose from three “emoji” faces representing neutral, “a little funny”, and funny. A trigger pulse from the scanner synchronized the onset of the experiment to the beginning of the acquisition of a functional MRI volume. Participants were instructed that they should wait for the movies to start. A technical problem meant that the first 61 participants had their experiment triggered one TR-time (2.25 s) earlier than the final 24 participants. This was corrected in the individual-level functional MRI analysis. All subjects were questioned after the scanning about the occurrence of cataplexy.

41 patients and 44 first-degree relatives rated the movies, but one patient and one first-degree relative could not be included in the ratings analysis because they rated all movies as neutral, confirming after the session that they did not think any of the movies were funny. All three rating categories were considered for the analysis of behavioral responses, but the ratings of “a little funny” and funny were combined to give a category (“fun”) for the functional MRI analysis. Eight participants had to stop the experiment and run it again later, mainly due to feeling drowsy/falling asleep under the first experiment. Due to a technical problem one participant watched the movies in black and white, and, due to human error, two participants viewed the movies without sound.

### MRI acquisition and processing

Imaging was conducted on a General Electric Discovery MR750 3T scanner at Oslo University Hospital using a 32-channel head coil. For registration purposes we acquired a T1-weighted scan (duration: 4 min 43 s) with voxel size 1 x 1 x 1 mm; repetition time (TR): 8.16 ms; echo time (TE): 3.18 ms; flip angle: 12°; 188 sagittal slices. We acquired functional MRI data with a T2*-weighted echo-planar imaging sequence (duration: 16 min 19 s) with 430 volumes; 3 mm slice thickness, in-plane resolution: 2.67 x 2.67; TR: 2250 ms; TE: 30 ms; slice gap: 0.5 mm; flip angle: 79°; 43 axial slices.

Functional MRI data were processed and analyzed with the FMRI Expert Analysis Tool (FEAT) from the Functional Magnetic Resonance Imaging of the Brain (FMRIB) Software Library (FSL) (https://fsl.fmrib.ox.ac.uk/fsl/fslwiki) (Smith *et al*., 2004; Jenkinson *et al*., 2012). Individual first-level preprocessing included motion correction with MCFLIRT (Jenkinson *et al*., 2002), spatial smoothing (full width at a half-maximum of 5 mm), grand-mean intensity normalization of the whole 4D dataset by a single scaling factor and high-pass temporal filtering (128 s).

We processed T1-weighted data in FreeSurfer (http://surfer.nmr.mgh.harvard.edu) to obtain brain-extracted T1-weighted volumes. The T1 images were further registered to the functional images using FMRIB's Linear Image Registration Tool (FLIRT) (Jenkinson and Smith, 2001; Jenkinson *et al*., 2002), optimized by boundary-based registration (Greve and Fischl, 2009), and then nonlinear registration to a standard MNI space was done using FMRIB’s Non-linear Image Registration Tool (FNIRT) (Andersson *et al*., 2007a; Andersson *et al*., 2007b).

### Statistical analysis

Individual-level general linear models (GLM) were fitted using FILM (FMRIB’s Improved Linear Model) modeling the movies (fun/neutral) and rating/response periods as blocks and the interspersed fixation periods as implicit baselines. The design matrix included nuisance regressors for six motion parameters and their derivatives. Temporal derivatives were added to account for regional differences in the timing of the hemodynamic response, e.g., due to differences in acquisition time between slices. Regressors were filtered and convolved with a double-gamma hemodynamic response function before the model fit. Single-subject contrasts were calculated for fun+, fun-, neutral+, neutral-, fun > neutral, neutral > fun, response+, response-. An additional analysis was performed to compare brain activation during movies without a humorous punchline (89.0% neutral-rated).

Contrast parameter estimates (COPEs) for each first-level contrast were concatenated in standard space and submitted to group analysis in Permutation Analysis of Linear Models (PALM) (Winkler *et al*., 2014; Winkler *et al*., 2015), testing for group differences between narcolepsy type 1 patients and first-degree relatives, while controlling for age and gender. We corrected for multiple testing by running 5000 permutations and threshold-free cluster enhancement (TFCE) as implemented in Permutation Analysis of Linear Models (PALM). To control for lack of independence (patients and first-degree relatives from the same family, two related patients, and siblings within the first-degree relative group) permutations were constrained between first-degree relatives. Corrected two-tailed values of *p* < 0.05 were considered significant. We also performed a separate analysis for a reduced sample (22 narcolepsy type 1 patients (15 females, age 21.5 ± 8.2 years) and 26 controls (13 females, age 20.6 ± 9.4 years), in which we excluded all patients and first-degree relatives with comorbidities, as well as all participants who had to re-watch the movies (*n* = 8) or had watched the movies in black and white (*n* = 1) or without sound (*n* = 2), and first-degree relatives who had experienced cataplexy-like episodes (*n* = 6). To assess the similarities in the results between the full and reduced samples, spatial correlations were computed between the uncorrected t-statistic maps.

For the behavioral responses we coded no rating, neutral, “a little funny” and funny as 0, 1, 2 and 3, respectively. We tested for group differences in average movie ratings (for all movies/only rated movies) using Permutation Analysis of Linear Models (PALM) (Winkler *et al*., 2014; Winkler *et al*., 2015), controlling for age, gender and the number of movies not rated. We ran 5000 permutations and, to control for lack of independence, these were constrained between first-degree relatives. Corrected two-tailed values of *p* < 0.05 were considered significant. We performed chi-square tests of independence to test for the relation between group and rating (neutral, “a little funny” and funny), including and excluding the movies without rating.

### Data availability

The data are not publicly available due to ethical restrictions as it could compromise the privacy of research participants.

## Results

### Behavioral responses

One patient reported a cataplectic attack during the scanning, but could not precisely identify during which movie it had occurred. Patients rated an average of (mean± SD) 11.5 ± 5.3 movies as neutral, 14.3 ± 6.6 movies as fun, and 4.2 ± 6.1 movies had no rating. Controls rated 13.7 ± 6.1 movies as neutral, 15.6 ± 6.3 movies as fun, and 0.8 ± 1.9 movies had no rating. Considering only rated trials, patients and controls rated an average of 55.5% and 53.2% of movies as fun, respectively. Permutation testing revealed no significant group difference in average movie ratings when considering all movies or only rated movies. The chi square test of independence was not significant when excluding the movies without rating, but it was significant when including these movies.

### Brain activation

#### Main effects and group differences for fun-rated movies in the full and reduced samples

Figure 1 summarize the results for fun-rated movies in the full sample. Both groups showed significant activations in the bilateral temporal poles, tempo-parietal areas, temporo-occipito-parietal areas, amygdala, hippocampus, thalamus, occipital areas, and in the right inferior frontal gyrus. Permutation testing revealed no significant group differences between patients and controls. Subsequent sub-analysis of the reduced sample (excluding participants with comorbidity, first-degree relatives who had experienced cataplexy-like episodes, participants who had to re-watch the movies, or watch the movies in black and white or without sound) yielded similar results for the main effects of fun, and no significant group differences.

**Figure 1.**
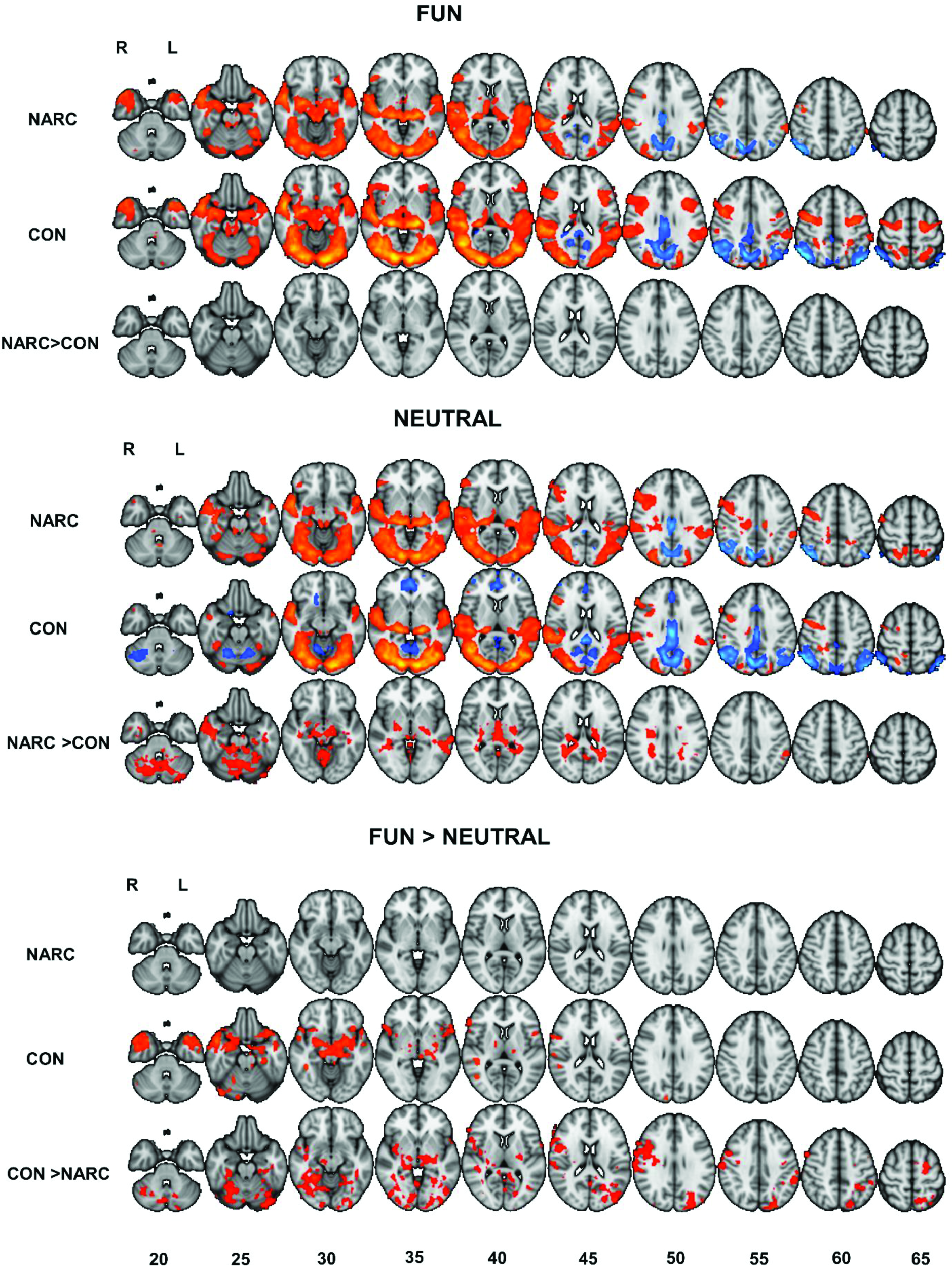
Main effects of task on brain activation and group comparisons. Summarizing functional MRI results with main effects and group comparisons. Numbers reflect the z-coordinate in the MNI 2-mm space. Only voxels with a two-tailed value of p < 0.05, corrected for multiple comparisons using permutation testing and TFCE (threshold-free cluster enhancement) are shown. Narc: narcolepsy type 1 patients, Con: first-degree relatives (controls). R: right, L: left. Red/Orange: higher activation. Blue: lower activation.

#### Main effects and group differences for neutral-rated movies in the full and reduced samples

Figures 1 and 2 summarize the results for neutral-rated movies in the full sample. Both groups showed significant activation in the bilateral thalamus, occipital areas, temporal areas and right inferior frontal gyrus. Patients showed significantly higher activation for neutral-rated movies compared with controls in widespread areas (16 267 voxels), including bilaterally in the thalamus, pallidum, putamen, amygdala, hippocampus, middle temporal gyrus, cerebellum, brainstem and in the left precuneus, supramarginal gyrus and caudate.

**Figure 2.**
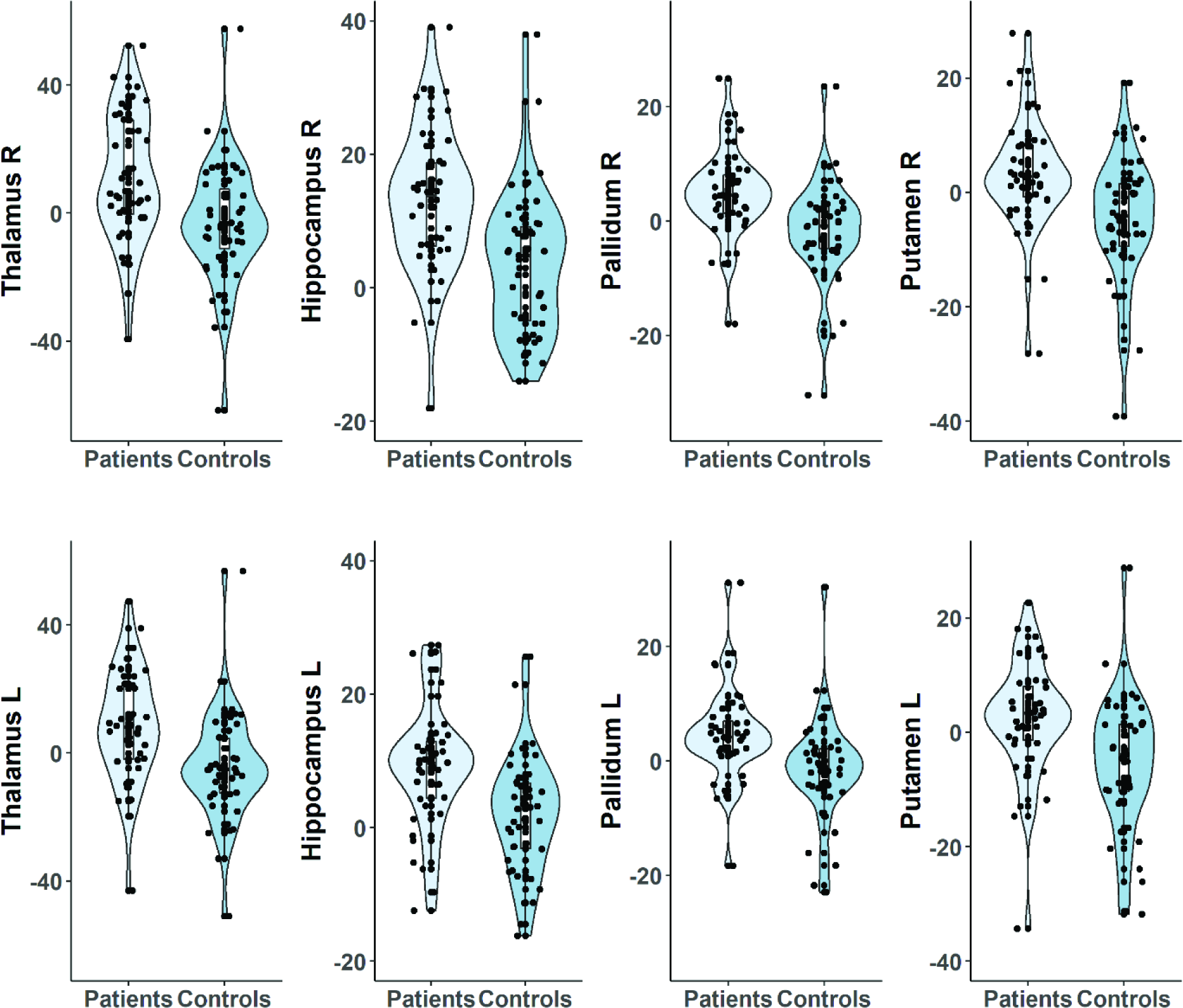
Group-wise distributions of COPE values for neutral+ across select regions of interest. The violin plots show the mean COPEs in regions of interest with significant group differences (identified by permutation testing) in the first-level contrast neutral+. The probabilistic atlas, Harvard-Oxford Subcortical Structural Atlas, implemented in FSL (https://fsl.fmrib.ox.ac.uk/fsl/fslwiki) (Smith *et al*., 2004; Jenkinson *et al*., 2012) was used to extract masks (threshold = 10) of regions of interests. Patients: Narcolepsy type 1 patients, Controls: first-degree relatives. R: right, L: left

Sub-analysis of the reduced sample (excluding participants with comorbidity, first-degree relatives who had experienced cataplexy-like episodes, participants who had to re-watch the movies, or watch the movies in black and white or without sound) revealed even more widespread group differences than in the full sample (44 538 voxels), with many areas exhibiting significantly higher activations in patients compared to controls, including bilaterally in amygdala, thalamus, putamen, hippocampus, caudate, pallidum, insula, paracingulate gyrus, cingulate gyrus, middle temporal gyrus, precuneus, precentral gyrus, inferior frontal gyrus, supramarginal gyrus, cerebellum, frontal areas, temporal areas and brainstem (Supplementary Figure 1). The spatial correlation between the uncorrected t-statistic maps obtained from the group comparisons in the full and reduced samples was 0.83, suggesting a highly similar pattern and direction of effects.

#### Main effects and group differences between fun and neutral in full and reduced samples

Figures 1 and 3 summarize the results for the fun *vs*. neutral contrast in the full sample. In controls, several areas showed significantly higher activation for fun-rated compared with neutral-rated movies. These included the bilateral temporal poles, amygdala, inferior frontal gyrus, thalamus, putamen and frontalorbital cortex. Conversely, patients showed no significant differences in brain activation between fun and neutral-rated movies.

**Figure 3.**
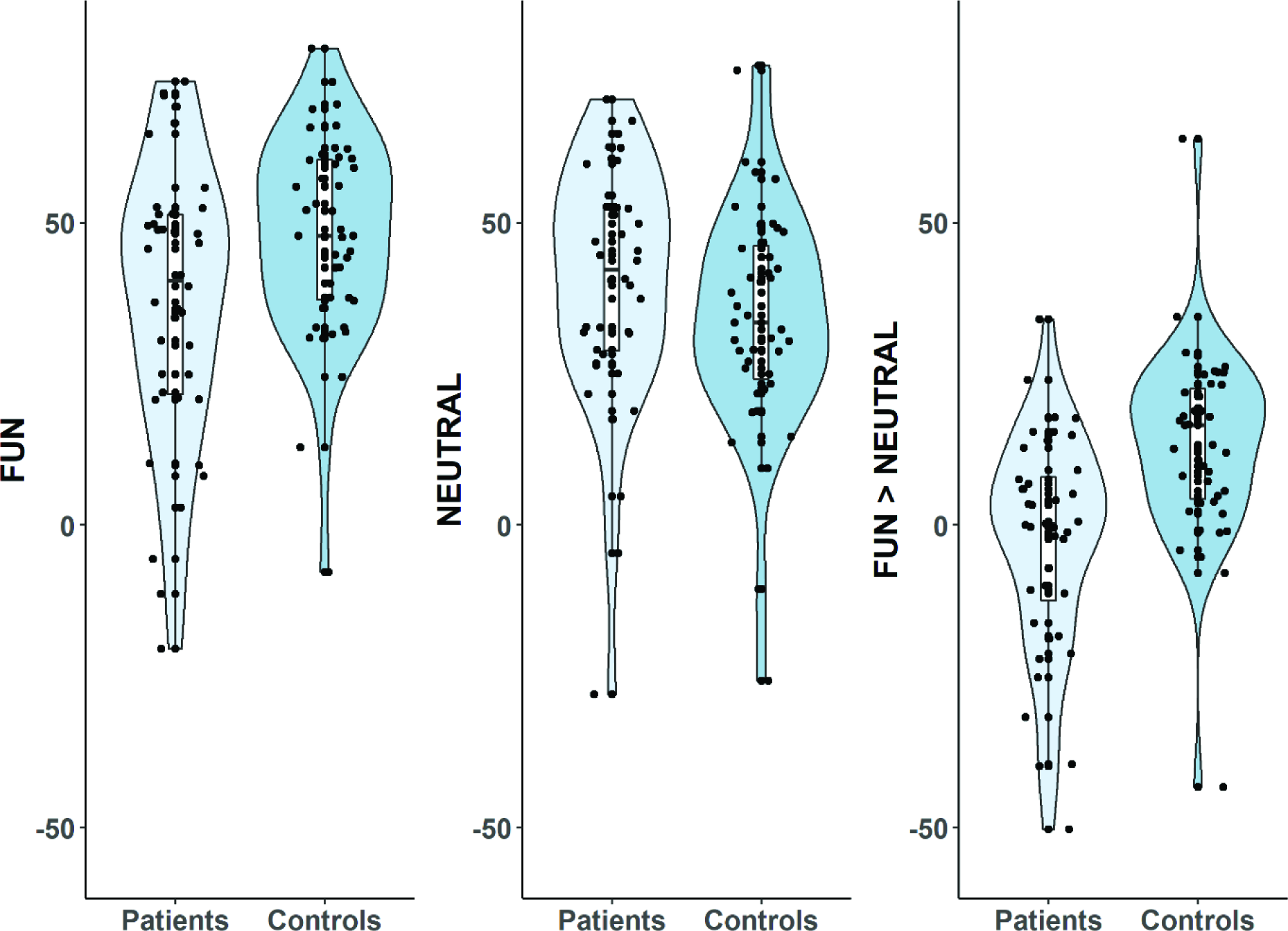
Group-wise distributions of COPE values across all significant voxels. The violin plots show the mean COPEs for the first-level contrasts; fun+, neutral+, fun > neutral, for voxels (16 604 voxels) with a significant group difference in fun > neutral identified by permutation testing. Patients: Narcolepsy type 1 patients, Controls: first-degree relatives.

Hence, group comparisons for fun > neutral revealed several areas with a significantly higher increase in activation for controls compared with patients (16 604 voxels), including bilaterally in the inferior frontal gyrus, thalamus, putamen, precentral gyrus, lingual gyrus, supramarginal gyrus, occipital areas, temporal areas, cerebellum, and in the right hippocampus, postcentral gyrus, pallidum and insula.

Group comparison in the reduced sample (excluding participants with comorbidity, first-degree relatives who had experienced cataplexy-like episodes, participants who had to re-watch the movies, or watch the movies in black and white or without sound) revealed an even more widespread (28 471 voxels) pattern of significantly higher activation in fun > neutral in controls compared with patients, largely overlapping with the previously described areas for the full sample, but also including bilaterally the hippocampus, the right amygdala and left caudate. The spatial correlation between the uncorrected t-statistic map was 0.76, suggesting a highly similar pattern and direction of effects.

#### Main effect and group differences in movies without a humorous punchline

Figure 4 summarizes the main effect for movies without a humorous punchline (89.0% neutral-rated). Similar brain activations were seen in patients and controls, with activations in several areas, including bilaterally in the thalamus, temporal and occipital areas. We found no significant group differences in activations for movies without a humorous punchline.

**Figure 4.**
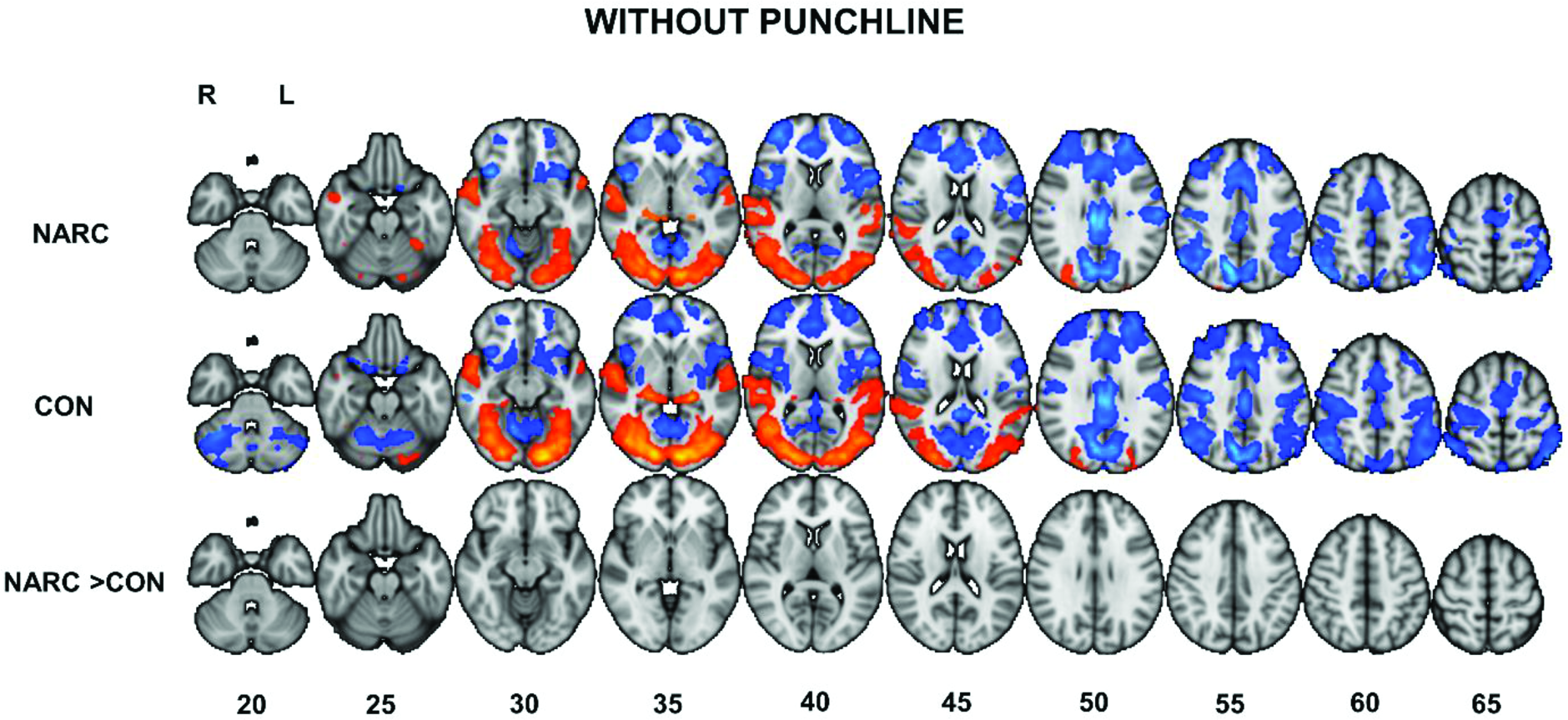
Functional MRI results for movies without a humorous punchline. Summarizing functional MRI results with main effects and group comparisons for movies without a humorous punchline. There were no significant differences in activations between narcolepsy type 1 patients and controls. Numbers reflect the z-coordinate in MNI 2-mm space. Only voxels with a two-tailed value of p < 0.05, corrected for multiple comparisons using permutation testing and TFCE (threshold-free cluster enhancement), are shown. Narc: narcolepsy type 1 patients, Con: first-degree relatives (controls). R: right, L: left. Red/Orange: higher activation. Blue: lower activation.

## Discussion

Theoretical and empirical work has suggested that abnormal humor processing may take place in the brain during fun stimuli and cataplexy in narcolepsy type 1 patients, but previous studies with limited sample sizes (Reiss *et al*., 2008; Schwartz *et al*., 2008; Meletti *et al*., 2015) have not provided a coherent account of the brain activations patterns. Further, they did not explore the brain activation patterns of other parts of humor processing with neutral-rated movies, which might contain a humorous punchline and therefore have the potential to be experienced as funny, despite being rated as neutral, or movies without a humorous punchline.

The work presented here is the largest functional MRI study of humor processing in narcolepsy type 1 patients to date, and the first to examine well-characterized post-H1N1 narcolepsy type 1 patients. We report significantly higher brain activation to neutral-rated movies in narcolepsy type 1 patients compared to controls. This activation involves several regions, including bilaterally in the thalamus, pallidum, putamen, amygdala, hippocampus, middle temporal gyrus, cerebellum, brainstem and in the left precuneus, supramarginal gyrus and caudate. We found no significant differences in brain activation between patients and controls for fun-rated movies. In accordance with these observations, there was no significant difference in brain activation in patients between fun-rated and neutral-rated movies, in contrast to controls where several regions showed significantly stronger differentiation between fun-rated and neutral-rated movies compared with patients including bilaterally in the inferior frontal gyrus, thalamus, putamen, precentral gyrus, lingual gyrus, supramarginal gyrus, occipital areas, temporal areas, cerebellum, and in the right hippocampus, postcentral gyrus, pallidum and insula.

Three previous functional MRI studies have studied humor processing in narcolepsy type 1 patients using humorous pictures (Reiss *et al*., 2008; Schwartz *et al*., 2008) or movies (Meletti *et al*., 2015). Cataplexy attacks were elicited in zero (Schwartz *et al*., 2008), one (Reiss *et al*., 2008) and 10 (Meletti *et al*., 2015) patients. The first study (Schwartz *et al*., 2008) compared 12 narcolepsy type 1 patients (all with cataplexy and unmedicated for at least 14 days before the scanning) with 12 healthy controls. Eight patients had reduced or undetectable levels of CSF-hypocretin. The authors reported that patients and controls rated similar proportions of images as funny. Patients compared to controls had lower activation in several regions including the hypothalamus, and higher activation in several regions including the amygdala, to humorous compared to neutral pictures.

The second study (Reiss *et al*., 2008) included 10 narcolepsy type 1 patients, all with cataplexy and who were unmedicated for at least 5 days before scanning, and 10 healthy controls. Six patients had low levels of hypocretin. Unlike the first study (Schwartz *et al*., 2008), the authors reported that patients rated significantly fewer cartoons as funny compared with the controls. Further they reported that patients showed increased brain activation compared to controls in several regions including the hypothalamus, the ventral striatum and right inferior frontal gyrus when looking at funny compared with non-funny cartoons (pictures).

The third study (Meletti *et al*., 2015) took a different approach by focusing on eliciting cataplectic attacks in the scanner. 21 narcolepsy type 1 patients (all, drug-naïve, hypocretin-deficient and with cataplexy) were studied with functional MRI acquired with synchronously EEG, while watching funny movies that were tailored to each patient’s preference. The study did not include any healthy controls. 10 patients had cataplectic attacks and 16 experienced laughter. Laughter was associated with an increased brain response bilaterally in the anterior cingulate gyrus and the motor/premotor cortex. Cataplexy was associated with increased brain response in several areas, including; the amygdala, anterior insular cortex, ventromedial prefrontal cortex, nucleus accumbens, locus coeruleus and the anteromedial pons.

In our study, we found that narcolepsy type 1 patients showed no differences from the control group in their average ratings of the movies, similar to the findings of one study (Schwartz *et al*., 2008) but in contrast to those of another (Reiss *et al*., 2008). Further, we found that several brain regions previously considered associated with humor processing (Vrticka *et al*., 2013) were activated in both patients and controls during fun-rated movies, but, in contrast to the other studies (Reiss *et al*., 2008; Schwartz *et al*., 2008), we found no group differences in brain activations for fun-rated movies. However, interestingly, there were several brain regions with higher activation during neutral-rated movies in patients compared with controls. Importantly, the neutral-rated movies in our study could have a potentially humorous punchline even if they were not subjectively rated as fun. A sub-analysis of five of the 30 movies without a humorous punchline revealed no significant differences in activations between patients and controls. The movies without a humorous punchline (89.0% neutral-rated) are otherwise similar in build-up and the anticipation that something funny might happen (even though it does not), which suggests that the recognition of a humorous punchline plays an important role in abnormal humor processing in narcolepsy type 1.

In short, we found that the narcolepsy type 1 patients compared to controls have a normal ability to subjectively rate humorous and neutral movies, but that neutral-rated movies with a potential humorous punchline elicit a brain overactivation in patients that is similar to the activation observed in response to fun-rated movies.

Due to our overall focus on the mechanisms of cataplexy, we were particularly interested in brain regions that have been linked to humor/cataplexy and REM sleep. One theory suggests that cataplexy represents dissociated REM sleep appearing while awake (Saper *et al*., 2010; Dauvilliers *et al*., 2014; Kornum *et al*., 2017). An alternative theory suggests that cataplexy is a variant of tonic immobility, which can be seen in animals that are unable to move in situations they experience to be dangerous (Overeem *et al*., 2002). However, this type of tonic immobility has not been observed in humans or other primates, and the most potent trigger for cataplexy is usually strong, positive emotions (thinking of, hearing, or telling a joke) (Overeem *et al*., 2002; Kornum *et al*., 2017).

Interestingly, in the present study, during neutral-rated movies, the thalamus showed significantly higher activation bilaterally in patients compared with controls. The thalamus has previously been shown to activate in response to humor in several studies (Mobbs *et al*., 2003; Wild *et al*., 2006; Kohn *et al*., 2011; Vrticka *et al*., 2013). Further, the thalamus has repeatedly been shown to have higher activation for REM sleep in functional MRI (Wehrle *et al*., 2005; Wehrle *et al*., 2007; Miyauchi *et al*., 2009) and PET (Maquet *et al*., 1996; Braun *et al*., 1997; Maquet and Franck, 1997; Buchsbaum *et al*., 2001) studies with polysomnographic monitoring.

Putamen, pallidum and caudate also showed significant overactivation in our patients compared with controls during neutral-rated movies, and have also been associated with REM sleep and humor. The basal ganglia show increased activity in PET (Braun *et al*., 1997) and functional MRI (Wehrle *et al*., 2005; Miyauchi *et al*., 2009) studies with polysomnography recordings during REM sleep. Humor perception has been found to correlate with higher activation in the pallidum and putamen (Wild *et al*., 2006), amusing films have been associated with higher activation in the right putamen and left globus pallidus (Goldin *et al*., 2005), and a meta-analysis reported that 70% of happiness-induction studies reported activation in the basal ganglia (Phan *et al*., 2002).

In the present study, we have used first-degree relatives as controls, and previous studies (Mignot, 1998; Ohayon *et al*., 2005; Wing *et al*., 2011) have reported a higher risk of developing narcolepsy in first-degree relatives of narcolepsy patients. However, all controls in our study were objectively ICSD-3-evaluated for a DG47.4 narcolepsy diagnosis, which was excluded in all controls. Six controls had experienced signs of cataplexy-like muscle weakness and although this was rare, the episodes had been provoked by emotional triggers like laughter, fun or excitement, and surprise. These controls were included in the full sample, but excluded from the sub-analysis of the reduced sample. Additionally, in the reduced sample we excluded patients and first-degree relatives with comorbidities and the participants who had to re-watch the movies or had watched the movies in black and white or without sound.

The results in the reduced sample were similar to those from the full sample, but even more widespread. There were more voxels showing significantly higher activation in patients compared with controls for neutral-rated movies in the reduced sample (44 538 voxels) compared with the full sample (16 267 voxels). The similarity of the findings in the full and reduced samples was further supported by the spatial correlation between the uncorrected test statistics suggesting very similar patterns and direction of effects. Likewise, there were similar group differences in the full and reduced sample when comparing fun and neutral-rated movies, but more widespread in the reduced sample. We observed significantly higher activation in controls compared with patients, in more voxels in the reduced sample (28 471 voxels) than in the full sample (16 604 voxels). Again, the spatial correlation between the test statistics indicated a very similar pattern and direction of effects in the full and reduced samples. Since the reduced sample have similar, but even more widespread results, the inclusion of participants with comorbidities, first-degree relatives with cataplexy-like episodes, participants who re-watched the movies or had watched movies in black and white or without sound, might have diminished some of the results in the full sample. However, it is a strength that the results in full and reduced samples support the same conclusions.

Mainly due to drowsiness/falling asleep, eight participants had to stop the experiment and restart it later. These were excluded from the reduced sample sub-analysis. Re-watching movies can give reduced activation in several regions, including the amygdala (Jaaskelainen *et al*., 2016), and we did find significantly higher activity in the amygdala for the reduced sample in patients compared with controls for neutral-rated movies. When comparing fun and neutral-rated movies in the reduced sample, controls showed significantly higher brain activation in the right amygdala compared with patients. The amygdala has been reliably associated with humor appreciation (Vrticka *et al*., 2013). Interestingly, PET (Maquet *et al*., 1996; Maquet and Franck, 1997; Nofzinger *et al*., 1997), and functional MRI (Miyauchi *et al*., 2009) studies have shown amygdala activation during REM sleep in healthy individuals. Further, electrophysiological studies have revealed changes in activity in the amygdala of narcoleptic dogs during cataplexy (Gulyani *et al*., 2002), amygdala lesions have been shown to reduce cataplexy in hypocretin knock-out mice (Burgess *et al*., 2013), and increased amygdala activity occurred during cataplexy in humans (Meletti *et al*., 2015), all of which suggest a role for the amygdala in mechanisms eliciting cataplexy.

Hippocampus activation is positively associated with a subjective rating of funniness (Iidaka, 2017), humor-induced smiling (Wild *et al*., 2006) and have also shown higher activation during REM-sleep in a PET study (Braun *et al*., 1997). We found that the hippocampus, like the amygdala, showed significantly higher activation in patients during neutral-rated movies compared with controls (in the full and reduced samples).

In our study, 92.7% of the patients had been H1N1-vaccinated, although the time of disease onset was changed to before the H1N1-vaccinations for three patients after a thorough evaluation of their medical history and records. Few differences have so far been found between sporadic and H1N1-vaccinated narcolepsy, except for a higher frequency of disturbed nocturnal sleep, shorter mean sleep latency (Pizza *et al*., 2014), a sudden onset of symptoms (Partinen *et al*., 2012; Heier *et al*., 2013), and more frequently sleep-onset REM periods (Dauvilliers *et al*., 2013). Post-H1N1 influenza narcolepsy also shows novel genetic associations (Han *et al*., 2013), but so far it seems unlikely that cataplexy mechanisms in H1N1-vaccinated narcolepsy are different from cataplexy in sporadic narcolepsy.

We included two patients with hypocretin deficiency, but without cataplexy, as prospective studies show that a substantial proportion of hypocretin-deficient non-cataplectic patients will later develop cataplexy (Andlauer *et al*., 2012).

In conclusion, we report functional MRI-based evidence of abnormal neuronal humor-processing in narcolepsy type 1 patients compared with controls, particularly an overactivation in response to neutral-rated movies in regions previously shown to be associated with humor, cataplexy and REM-sleep, including the thalamus, amygdala, hippocampus and basal ganglia. Unlike controls, patients showed similar activation during neutral-rated and fun-rated movies, so there is no significant differentiation of functional MRI activations between these two states, which might provide insight into the mechanisms associated with cataplexy. This “overactive” brain state during neutral-rated movies (but with a potentially humorous punchline) might represent a risk (hypersensitivity to potential humorous stimuli) for the narcolepsy type 1 patients which seem to have a lower threshold for activating the humor response (even when they subjectively rate the movie as neutral).

## Acknowledgements

We thank Phil Mason for editing the English text.

## Supplementary material

**Supplementary Figure 1.**
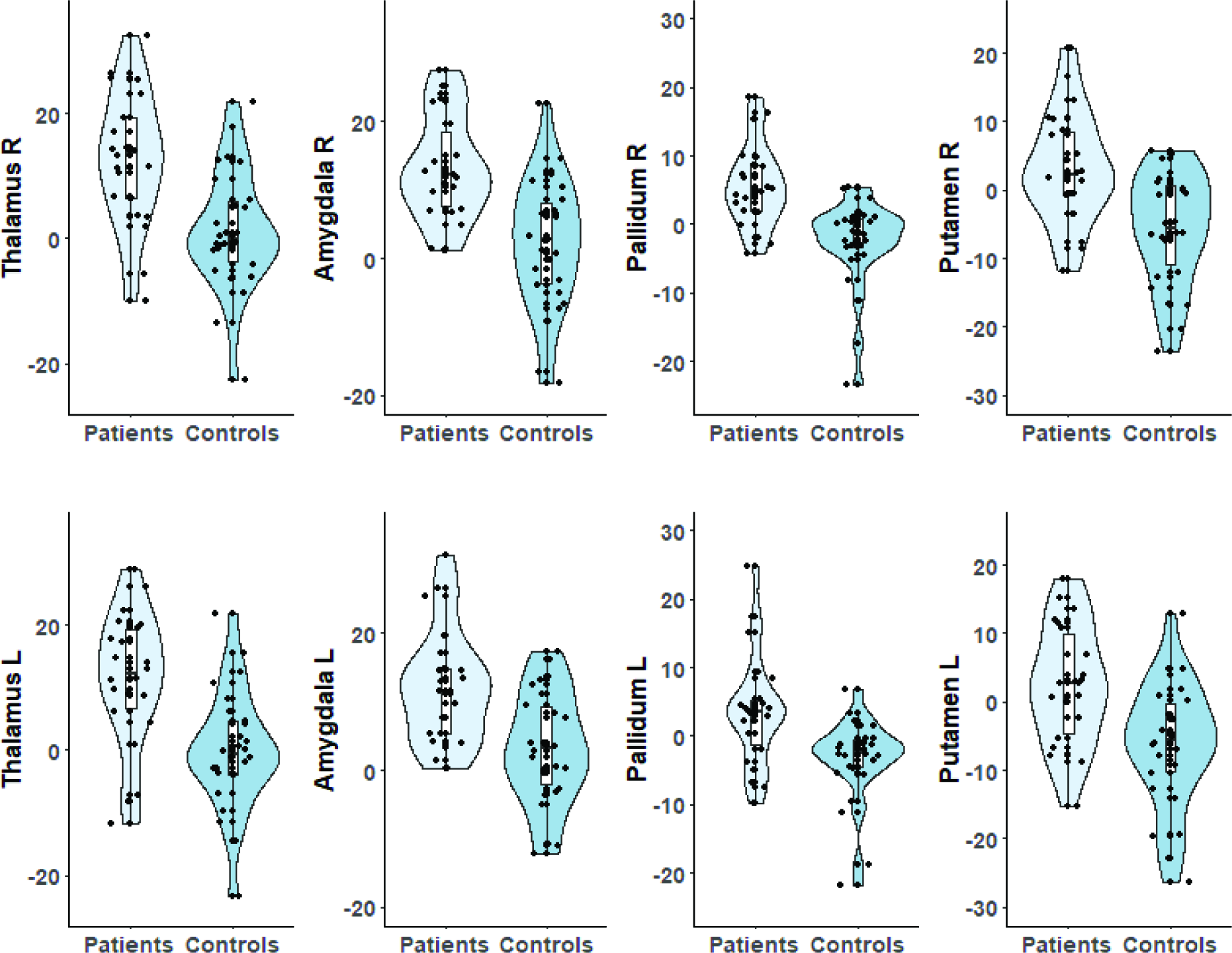
Mean contrast parameter estimates (COPEs) for regions of interest in neutral+ in the reduced sample. In the reduced sample of 22 narcolepsy type 1 patients (15 females, age 21.5 ± 8.2 years) and 26 controls (13 females, age 20.6 ± 9.4 years), we excluded all patients and first-degree relatives with comorbidity, as well as all participants who had to re-watch the movies or watch the movies in black and white or without sound, and first-degree relatives who had experienced cataplexy-like episodes. The violin plots show the mean COPEs for voxels with significant group differences (identified by permutation testing) in the first-level contrast neutral+ for regions of interest in the reduced sample. The probabilistic atlas Harvard-Oxford Subcortical Structural Atlas implemented in FSL (https://fsl.fmrib.ox.ac.uk/fsl/fslwiki) (Smith *et al*., 2004; Jenkinson *et al*., 2012) was used to extract masks (threshold = 10) of regions of interest. Patients: Narcolepsy type 1 patients, Controls: first-degree relatives. R: right, L: left

## Abbreviations

FSL: Functional Magnetic Resonance Imaging of the Brain Software Library
HLA: human leukocyte antigen
ICSD: International Classification of Sleep Disorders
MSLT: multiple sleep latency test
REM sleep: rapid eye movement sleep
TE: echo time
TR: repetition time

## References

American Academy of Sleep Medicine(AASM). International Classification of Sleep Disorders (ICSD) 3ed; 2014.

Andersson JLR, Jenkinson M, Smith S. Non-linear optimisation. FMRIB technical report TR07JA1 from http://www.fmrib.ox.ac.uk/datasets/techrep/tr07ja1/tr07ja1.pdf; 2007a.

Andersson JLR, Jenkinson M, Smith S. Non-linear registration, aka Spatial normalisation. FMRIB technical report TR07JA2 from http://www.fmrib.ox.ac.uk/datasets/techrep/tr07ja2/tr07ja2.pdf; 2007b.

Andlauer O, Moore Ht, Hong SC, Dauvilliers Y, Kanbayashi T, Nishino S, et al. Predictors of hypocretin (orexin) deficiency in narcolepsy without cataplexy. Sleep 2012; 35(9): 1247–55f.

Anic-Labat S, Guilleminault C, Kraemer HC, Meehan J, Arrigoni J, Mignot E. Validation of a cataplexy questionnaire in 983 sleep-disorders patients. Sleep 1999; 22(1): 77–87.

Braun AR, Balkin TJ, Wesenten NJ, Carson RE, Varga M, Baldwin P, et al. Regional cerebral blood flow throughout the sleep-wake cycle. An H2(15)O PET study. Brain : a journal of neurology 1997; 120 (Pt 7): 1173–97.

Buchsbaum MS, Hazlett EA, Wu J, Bunney WE, Jr. Positron emission tomography with deoxyglucose-F18 imaging of sleep. Neuropsychopharmacology : official publication of the American College of Neuropsychopharmacology 2001; 25(5 Suppl): S50–6.

Burgess CR, Oishi Y, Mochizuki T, Peever JH, Scammell TE. Amygdala lesions reduce cataplexy in orexin knock-out mice. The Journal of neuroscience : the official journal of the Society for Neuroscience 2013; 33(23): 9734–42.

Dauvilliers Y, Arnulf I, Lecendreux M, Monaca Charley C, Franco P, Drouot X, et al. Increased risk of narcolepsy in children and adults after pandemic H1N1 vaccination in France. Brain : a journal of neurology 2013; 136(Pt 8): 2486–96.

Dauvilliers Y, Arnulf I, Mignot E. Narcolepsy with cataplexy. Lancet (London, England) 2007; 369(9560): 499–511.

Dauvilliers Y, Siegel JM, Lopez R, Torontali ZA, Peever JH. Cataplexy--clinical aspects, pathophysiology and management strategy. Nature reviews Neurology 2014; 10(7): 386–95.

Goldin PR, Hutcherson CA, Ochsner KN, Glover GH, Gabrieli JD, Gross JJ. The neural bases of amusement and sadness: a comparison of block contrast and subject-specific emotion intensity regression approaches. NeuroImage 2005; 27(1): 26–36.

Greve DN, Fischl B. Accurate and robust brain image alignment using boundary-based registration. NeuroImage 2009; 48(1): 63–72.

Gulyani S, Wu MF, Nienhuis R, John J, Siegel JM. Cataplexy-related neurons in the amygdala of the narcoleptic dog. Neuroscience 2002; 112(2): 355–65.

Han F, Faraco J, Dong XS, Ollila HM, Lin L, Li J, et al. Genome wide analysis of narcolepsy in China implicates novel immune loci and reveals changes in association prior to versus after the 2009 H1N1 influenza pandemic. PLoS genetics 2013; 9(10): e1003880.

Heier MS, Evsiukova T, Vilming S, Gjerstad MD, Schrader H, Gautvik K. CSF hypocretin-1 levels and clinical profiles in narcolepsy and idiopathic CNS hypersomnia in Norway. Sleep 2007; 30(8): 969–73.

Heier MS, Gautvik KM, Wannag E, Bronder KH, Midtlyng E, Kamaleri Y, et al. Incidence of narcolepsy in Norwegian children and adolescents after vaccination against H1N1 influenza A. Sleep medicine 2013; 14(9): 867–71.

Iidaka T. Humor Appreciation Involves Parametric and Synchronized Activity in the Medial Prefrontal Cortex and Hippocampus. Cerebral cortex (New York, NY : 1991) 2017; 27(12): 5579–91.

Jaaskelainen IP, Pajula J, Tohka J, Lee HJ, Kuo WJ, Lin FH. Brain hemodynamic activity during viewing and re-viewing of comedy movies explained by experienced humor. Scientific reports 2016; 6: 27741.

Jenkinson M, Bannister P, Brady M, Smith S. Improved optimization for the robust and accurate linear registration and motion correction of brain images. NeuroImage 2002; 17(2): 825–41.

Jenkinson M, Beckmann CF, Behrens TE, Woolrich MW, Smith SM. FSL. NeuroImage 2012; 62(2): 782–90.

Jenkinson M, Smith S. A global optimisation method for robust affine registration of brain images. Medical image analysis 2001; 5(2): 143–56.

Juvodden HT, Alnaes D, Lund MJ, Agartz I, Andreassen OA, Dietrichs E, et al. Widespread white matter changes in post-H1N1 narcolepsy type 1 patients and 1st degree relatives. Sleep 2018.

Knudsen S, Jennum PJ, Alving J, Sheikh SP, Gammeltoft S. Validation of the ICSD-2 criteria for CSF hypocretin-1 measurements in the diagnosis of narcolepsy in the Danish population. Sleep 2010; 33(2): 169–76.

Kohn N, Kellermann T, Gur RC, Schneider F, Habel U. Gender differences in the neural correlates of humor processing: implications for different processing modes. Neuropsychologia 2011; 49(5): 888–97.

Kornum BR, Knudsen S, Ollila HM, Pizza F, Jennum PJ, Dauvilliers Y, et al. Narcolepsy. Nature reviews Disease primers 2017; 3: 16100.

Li J, Hu Z, de Lecea L. The hypocretins/orexins: integrators of multiple physiological functions. British journal of pharmacology 2014; 171(2): 332–50.

Mahler SV, Moorman DE, Smith RJ, James MH, Aston-Jones G. Motivational activation: a unifying hypothesis of orexin/hypocretin function. Nature neuroscience 2014; 17(10): 1298–303.

Maquet P, Franck G. REM sleep and amygdala. Molecular psychiatry 1997; 2(3): 195–6.

Maquet P, Peters J, Aerts J, Delfiore G, Degueldre C, Luxen A, et al. Functional neuroanatomy of human rapid-eye-movement sleep and dreaming. Nature 1996; 383(6596): 163–6.

Meletti S, Vaudano AE, Pizza F, Ruggieri A, Vandi S, Teggi A, et al. The Brain Correlates of Laugh and Cataplexy in Childhood Narcolepsy. The Journal of neuroscience : the official journal of the Society for Neuroscience 2015; 35(33): 11583–94.

Mignot E. Genetic and familial aspects of narcolepsy. Neurology 1998; 50(2 Suppl 1): S16–22.

Miyauchi S, Misaki M, Kan S, Fukunaga T, Koike T. Human brain activity time-locked to rapid eye movements during REM sleep. Experimental brain research 2009; 192(4): 657–67.

Mobbs D, Greicius MD, Abdel-Azim E, Menon V, Reiss AL. Humor modulates the mesolimbic reward centers. Neuron 2003; 40(5): 1041–8.

Nofzinger EA, Mintun MA, Wiseman M, Kupfer DJ, Moore RY. Forebrain activation in REM sleep: an FDG PET study. Brain research 1997; 770(1-2): 192–201.

Ohayon MM, Ferini-Strambi L, Plazzi G, Smirne S, Castronovo V. Frequency of narcolepsy symptoms and other sleep disorders in narcoleptic patients and their first-degree relatives. Journal of sleep research 2005; 14(4): 437–45.

Overeem S, Lammers GJ, van Dijk JG. Cataplexy: 'tonic immobility' rather than 'REM-sleep atonia'? Sleep medicine 2002; 3(6): 471–7.

Overeem S, van Nues SJ, van der Zande WL, Donjacour CE, van Mierlo P, Lammers GJ. The clinical features of cataplexy: a questionnaire study in narcolepsy patients with and without hypocretin-1 deficiency. Sleep medicine 2011; 12(1): 12–8.

Partinen M, Saarenpaa-Heikkila O, Ilveskoski I, Hublin C, Linna M, Olsen P, et al. Increased incidence and clinical picture of childhood narcolepsy following the 2009 H1N1 pandemic vaccination campaign in Finland. PloS one 2012; 7(3): e33723.

Peyron C, Faraco J, Rogers W, Ripley B, Overeem S, Charnay Y, et al. A mutation in a case of early onset narcolepsy and a generalized absence of hypocretin peptides in human narcoleptic brains. Nature medicine 2000; 6(9): 991–7.

Peyron C, Tighe DK, van den Pol AN, de Lecea L, Heller HC, Sutcliffe JG, et al. Neurons containing hypocretin (orexin) project to multiple neuronal systems. The Journal of neuroscience : the official journal of the Society for Neuroscience 1998; 18(23): 9996–10015.

Phan KL, Wager T, Taylor SF, Liberzon I. Functional neuroanatomy of emotion: a meta-analysis of emotion activation studies in PET and fMRI. NeuroImage 2002; 16(2): 331–48.

Pizza F, Peltola H, Sarkanen T, Moghadam KK, Plazzi G, Partinen M. Childhood narcolepsy with cataplexy: comparison between post-H1N1 vaccination and sporadic cases. Sleep medicine 2014; 15(2): 262–5.

Reiss AL, Hoeft F, Tenforde AS, Chen W, Mobbs D, Mignot EJ. Anomalous hypothalamic responses to humor in cataplexy. PloS one 2008; 3(5): e2225.

Saper CB. The neurobiology of sleep. Continuum (Minneapolis, Minn) 2013; 19(1 Sleep Disorders): 19–31.

Saper CB, Fuller PM, Pedersen NP, Lu J, Scammell TE. Sleep state switching. Neuron 2010; 68(6): 1023–42.

Schwartz S, Ponz A, Poryazova R, Werth E, Boesiger P, Khatami R, et al. Abnormal activity in hypothalamus and amygdala during humour processing in human narcolepsy with cataplexy. Brain : a journal of neurology 2008; 131(Pt 2): 514–22.

Smith SM, Jenkinson M, Woolrich MW, Beckmann CF, Behrens TE, Johansen-Berg H, et al. Advances in functional and structural MR image analysis and implementation as FSL. NeuroImage 2004; 23 Suppl 1: S208–19.

Thannickal TC, Moore RY, Nienhuis R, Ramanathan L, Gulyani S, Aldrich M, et al. Reduced number of hypocretin neurons in human narcolepsy. Neuron 2000; 27(3): 469–74.

Vrticka P, Black JM, Reiss AL. The neural basis of humour processing. Nature reviews Neuroscience 2013; 14(12): 860–8.

Wehrle R, Czisch M, Kaufmann C, Wetter TC, Holsboer F, Auer DP, et al. Rapid eye movement-related brain activation in human sleep: a functional magnetic resonance imaging study. Neuroreport 2005; 16(8): 853–7.

Wehrle R, Kaufmann C, Wetter TC, Holsboer F, Auer DP, Pollmacher T, et al. Functional microstates within human REM sleep: first evidence from fMRI of a thalamocortical network specific for phasic REM periods. The European journal of neuroscience 2007; 25(3): 863–71.

Wild B, Rodden FA, Rapp A, Erb M, Grodd W, Ruch W. Humor and smiling: cortical regions selective for cognitive, affective, and volitional components. Neurology 2006; 66(6): 887–93.

Wing YK, Chen L, Lam SP, Li AM, Tang NL, Ng MH, et al. Familial aggregation of narcolepsy. Sleep medicine 2011; 12(10): 947–51.

Winkler AM, Ridgway GR, Webster MA, Smith SM, Nichols TE. Permutation inference for the general linear model. NeuroImage 2014; 92: 381–97.

Winkler AM, Webster MA, Vidaurre D, Nichols TE, Smith SM. Multi-level block permutation. NeuroImage 2015; 123: 253–68.

